# Systems and classical biology approaches unraveled role of olfactory receptors in progression of kidney fibrosis

**DOI:** 10.1101/2020.07.29.225425

**Authors:** Ali Motahharynia, Shiva Moein, Farnoush Kiyanpour, Kobra Moradzadeh, Yousof Gheisari

**Author notes:** Corresponding Author: Shiva Moein, PhD. Regenerative Medicine Research Center, Isfahan University of Medical Sciences, Isfahan, 8174673461, Iran.

## Abstract

The olfactory receptors (ORs) which are mainly known as odor-sensors in the olfactory epithelium are distributed in several non-sensory tissues. Despite the specified role of some of these receptors in normal physiology of the kidney, little is known about their potential effect in renal disorders. In this study, using the holistic view of systems biology, it was determined that ORs are significantly changed during the progression of kidney fibrosis. For further validation, common differentially expressed ORs resulted from reanalysis of two time-course microarray datasets were selected for experimental evaluation in a validated murine model of unilateral ureteral obstruction. Transcriptional analysis demonstrated considerable changes in the expression pattern of *Olfr433, Olfr129, Olfr1393, and Olfr161* during the progression of kidney fibrosis. In conclusion, our results highlight the impact of systems biology in determination of the underlying mechanisms of chronic diseases and indicate the importance of time-course approaches to unravel the patterns of gene expression. The novel ORs proposed in this study could be the subject of further functional investigations in the kidney.

## Introduction

Olfactory receptors (ORs), belonging to a super-family of G protein-coupled receptors, are well-recognized for their role in odor-sensation in the olfactory epithelium (OE). This family of receptors was first discovered by Linda Buck and Richard Axel, leading to a Nobel prize in 2004 (Buck & Axel, 1991; Watts, 2004). More investigations on ORs determined that they are not only expressed in the OE but also non-sensory organs (Massberg & Hatt, 2018). The study by Parmentier et al. determined the functionality of these receptors in sperm chemotaxis during fertilization (Parmentier et al., 1992). Furthermore, it was demonstrated that these receptors have roles in cytoskeletal remodeling, pulmonary hyperplasia (An & Liggett, 2018), angiogenesis (Kim et al., 2015), as well as heart metabolism in the cardiovascular system (Jovancevic et al., 2017). Besides, identification of the role of these receptors in other parts of the body, such as skin (Busse et al., 2014; Gelis et al., 2016; Tsai et al., 2017), gastrointestinal tract (Braun, Voland, Kunz, Prinz, & Gratzl, 2007; Kalbe et al., 2016; C. Wu et al., 2015), and the immune system (Clark, Nurmukhambetova, Li, Munger, & Lees, 2016; Li et al., 2013) highlights the importance of further investigations on their function in the body.

One of the tissues, in which the presence and function of ORs is investigated, is the kidney (Halperin Kuhns et al., 2019; Miyamoto et al., 2016; Peti-Peterdi, Kishore, & Pluznick, 2016; Pluznick et al., 2009; Rajkumar, Aisenberg, Acres, Protzko, & Pluznick, 2014; Shepard & Pluznick, 2017). Studies using unsupervised high-throughput techniques have discovered the existence of these receptors in the kidney (Feldmesser et al., 2006; Flegel, Manteniotis, Osthold, Hatt, & Gisselmann, 2013). Nevertheless, few studies have specifically focused on their role in renal function. The study by Pluznick et al. determined the Olfr78 role in blood pressure regulation through interactions with the byproducts of gut microbiota (Pluznick et al., 2013). Furthermore, Shepard et al. demonstrated that Olfr1393 participates in glucose transportation in both normal and pathological states of the kidney (Shepard et al., 2016; Shepard, Koepsell, & Pluznick, 2019). Considering the magnitude of this gene family (around 1000 genes in the mouse and 400 genes in the human) as well as their role in chemo-sensation, more investigations are needed to uncover the role of other OR subtypes in kidney function and hemostasis (Halperin Kuhns et al., 2019; Shepard & Pluznick, 2016).

In order to understand key molecular mechanisms underlying the progression of kidney fibrosis, a time-course microarray dataset was reanalyzed. Studying the modular structure of the constructed network determined a subset of ORs with a significant change in expression. Likewise, this finding was additionally confirmed by analysis of another time-course dataset with similar traits. Experimental evaluation of common ORs between two datasets determined incremental expression patterns during progression of kidney fibrosis. These results provide an initial clue about the importance of ORs functions in the progression of kidney fibrosis.

## Results

In order to investigate the underlying molecular mechanisms activated during the progression of chronic kidney disease, two time-course microarray datasets, GSE36496 and GSE96571 were analyzed. GSE36496 contains transcriptomic data of the UUO and sham-operated C57BL/6 mice at days 1, 2, 5, and 9 post-operation. The results of this study by Wu *et al*. suggest CEBPB and HNF4A signaling pathways as important regulators of kidney fibrosis (B. Wu & Brooks, 2012). The other dataset, GSE96571 which is also deposited by Wu *et al*. comprises the UUO and sham samples at hours 0.5, 1, 3, 5, 7, 12 as well as days 1,3,5, and 7 post-operation. The single-time point analysis of this dataset by Wu *et al*. determined overexpression of stress responder genes in the first hours of obstruction besides nephrotoxic damage-related genes at later time points (B. Wu et al., 2018). This dataset was used for validation of our findings from the first dataset.

### Unsupervised evaluation of microarray dataset determined high quality of the fibrotic model

The quality assessment of the GSE36496 dataset by PCA demonstrated the separation of sham and UUO groups from each other. Furthermore, the UUO samples were separated based on the times of harvest, which determines the model quality (Figure 1a). This result was also validated by the hierarchical clustering (Figure 1b). The DEGs were determined using the LIMMA Package of R software, which according to our recent study, is the most reliable tool for the time-course analysis of microarray data (Moradzadeh, Moein, Nickaeen, & Gheisari, 2019). A Comparison of the UUO and sham samples determined 2583 DEGs with an adjusted p-value < 0 .05. The DEGs were used for the construction of a gene interaction network and further topological analysis (Figure 1c).

**Figure 1.**
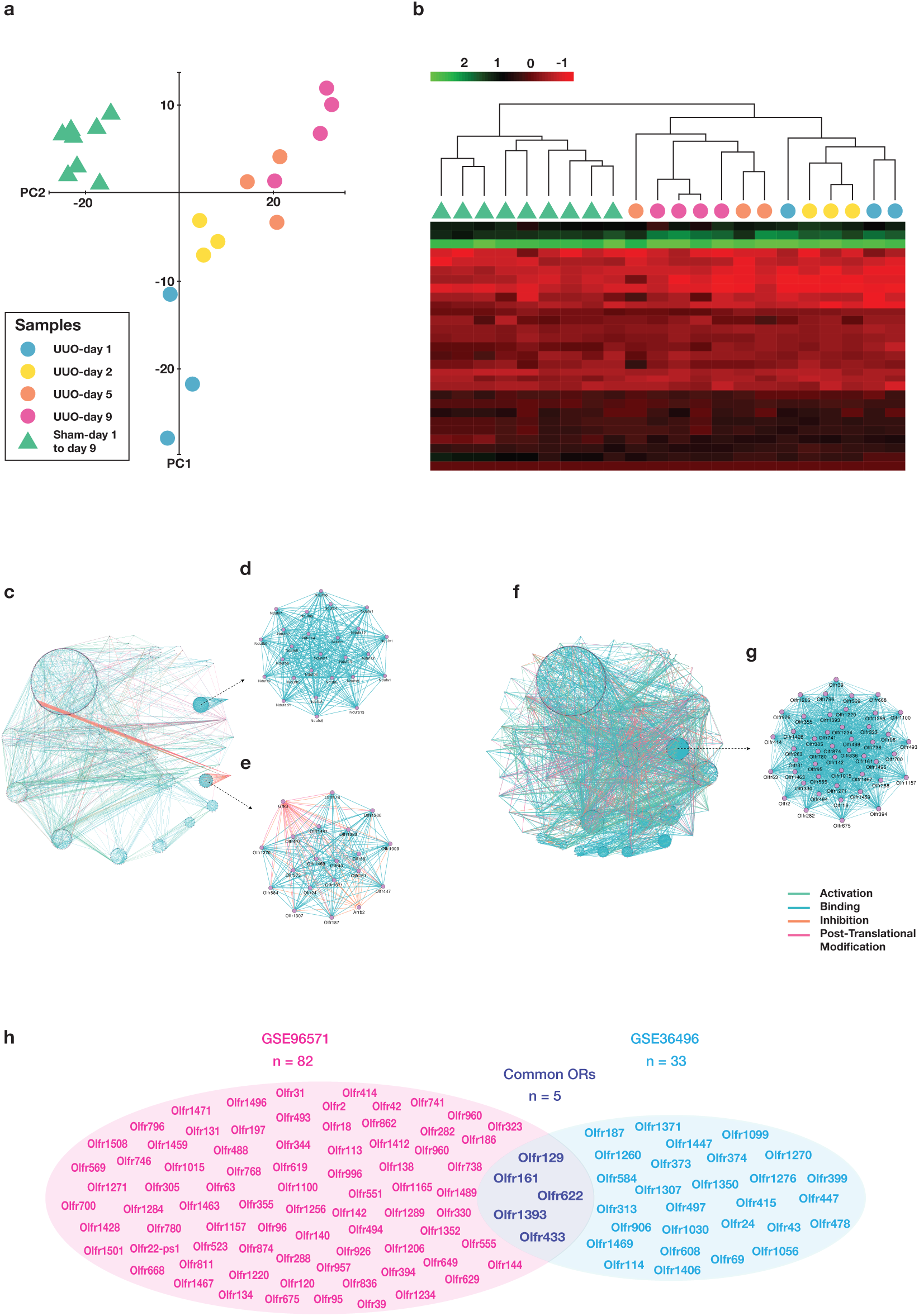
Network analysis of two microarray datasets on mouse model of ureteral obstruction. PCA analysis and heatmap clustering of GSE36496 dataset demonstrated good quality of samples regarding treatment and time of harvest (a, b). Gene interaction network created from 2583 DEGs of GSE36496 dataset (c). The first- (d) and the second-ranked (e) modules resulted from topological analysis of the network constitute NDUF and OR gene families, respectively. Gene Interaction network from 3176 DEGs of GSE96571 dataset (f). The top-ranked module from topological analysis of constructed network is constituted of Olfactory receptors (g). The overlap between differentially expressed ORs of GSE36496 and GSE96571 datasets (h).

### ORs constitute a dense module in the interactome map of kidney fibrosis

In order to discover the functional units of the network, module analysis was performed. Modules are individual units of biological network, which are similar in physical, chemical, or functional aspects and supposed to have a specific function in the networks (Choobdar et al., 2019; Lecca & Re, 2015). Assessment of densely connected regions of the network, based on the clustering coefficient, revealed 36 significant modules. The first and second modules were mainly related to Nduf and ORs signaling pathways (Figure 1d and e). NADH:ubiquinone oxidoreductase supernumerary subunits (NDUF) family of genes are expressed in the mitochondria and their relation to kidney fibrosis has previously been demonstrated by Granata *et al*. (Granata et al., 2009). The relationship between ORs and progression of chronic renal failure has also been investigated by only one study which determined the contribution of Olfr1393 in diabetes (Shepard et al., 2019).

For further investigation on the role of ORs in kidney fibrosis, another time-course UUO dataset was analyzed. LIMMA results determined 3176 DEGs (adjusted p-value < 0.05), which were further used for construction of a gene interaction network (Figure 1f). Topological analysis of the constructed network revealed 44 significant modules, of which ORs-related module obtained the first rank (Figure 1g). For validation of the results of microarray data analysis, common differentially expressed ORs (*Olfr433, Olfr129, Olfr1393, Olfr161, and Olfr622*) between two datasets were selected for in vivo expression analysis (figure 1h).

### Histopathological analysis of the UUO model validated the robustness of the constructed model

In order to experimentally evaluate the expression of selected ORs, mouse model of UUO was performed. Both UUO and sham-operated mice were followed over 21 days (Figure 2a). For validation of the quality of the constructed model, the histopathological analysis was done. Assessment of the sections revealed significant diffused tubulointerstitial fibrosis along with glomerular injuries, increased mesangial matrix, and diffused glomerulosclerosis in UUO-operated mice compared to the sham group over 21 days of UUO treatment (p-value < 0.05), all of which determined the robustness of constructed UUO model (Figure 2b and c).

**Figure 2.**
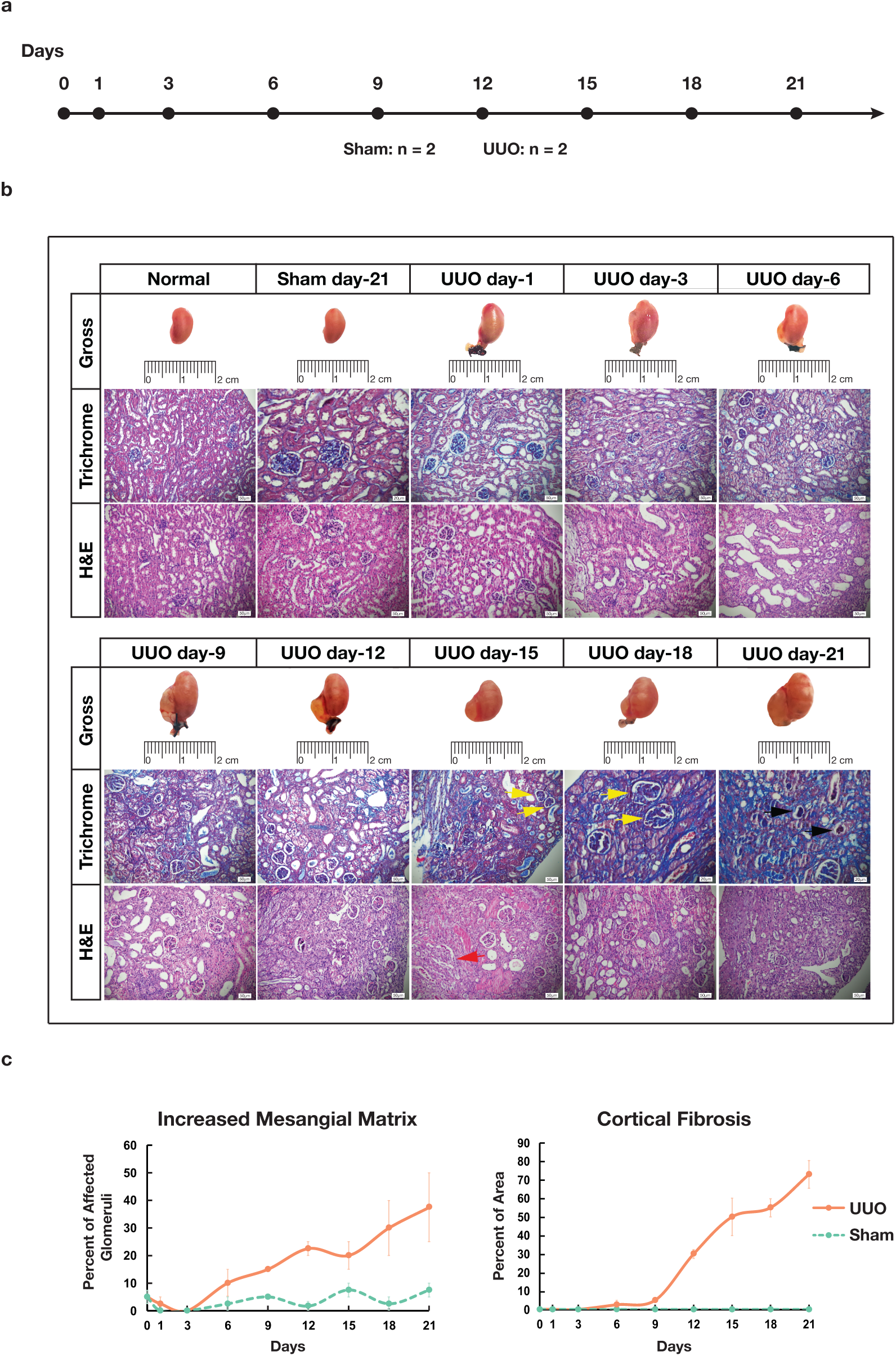
Histopathologic evaluation of UUO-operated mice over 21 days. The scheme of experimental design (a). Trichrome and H&E-stained renal sections in normal, sham (day-21), and UUO operated mice at days 1, 3, 6, 9, 12, 15, 18, and 21 post-surgery. Yellow, red, and black arrows stand for increased mesangial matrix, mesangial cell proliferation, and diffuse glomerulosclerosis, respectively (b). The percentages of glomeruli with increased mesangial matrix (p-value = 0) as well as cortical fibrosis (p-value = 3.25077e-10) for sham and UUO-operated mice. Data are mean ± SEM (c).

### Gene expression patterns demonstrated significant changes in ORs in the fibrotic kidney

As ORs belong to a highly conserved superfamily which most of them have a similar sequence, the specificity of designed primers was checked and validated by sanger sequencing (Supplementary data 1).

The expression of *Olfr433, Olfr129, Olfr1393, Olfr161, and Olfr622* was evaluated over 21 days in a time-course manner in both UUO and sham-operated mice (Figure 3). Except for *Olfr1393*, other genes demonstrated upregulation over time in comparison to the sham group. To test whether these changes between the sham and UUO group were significant during time, a two-way ANOVA analysis was applied. The results of the analysis determined significant changes in patterns of expression in the UUO group compared to sham for all the genes except for *Olfr622* (p-value < 0.05).

**Figure 3.**
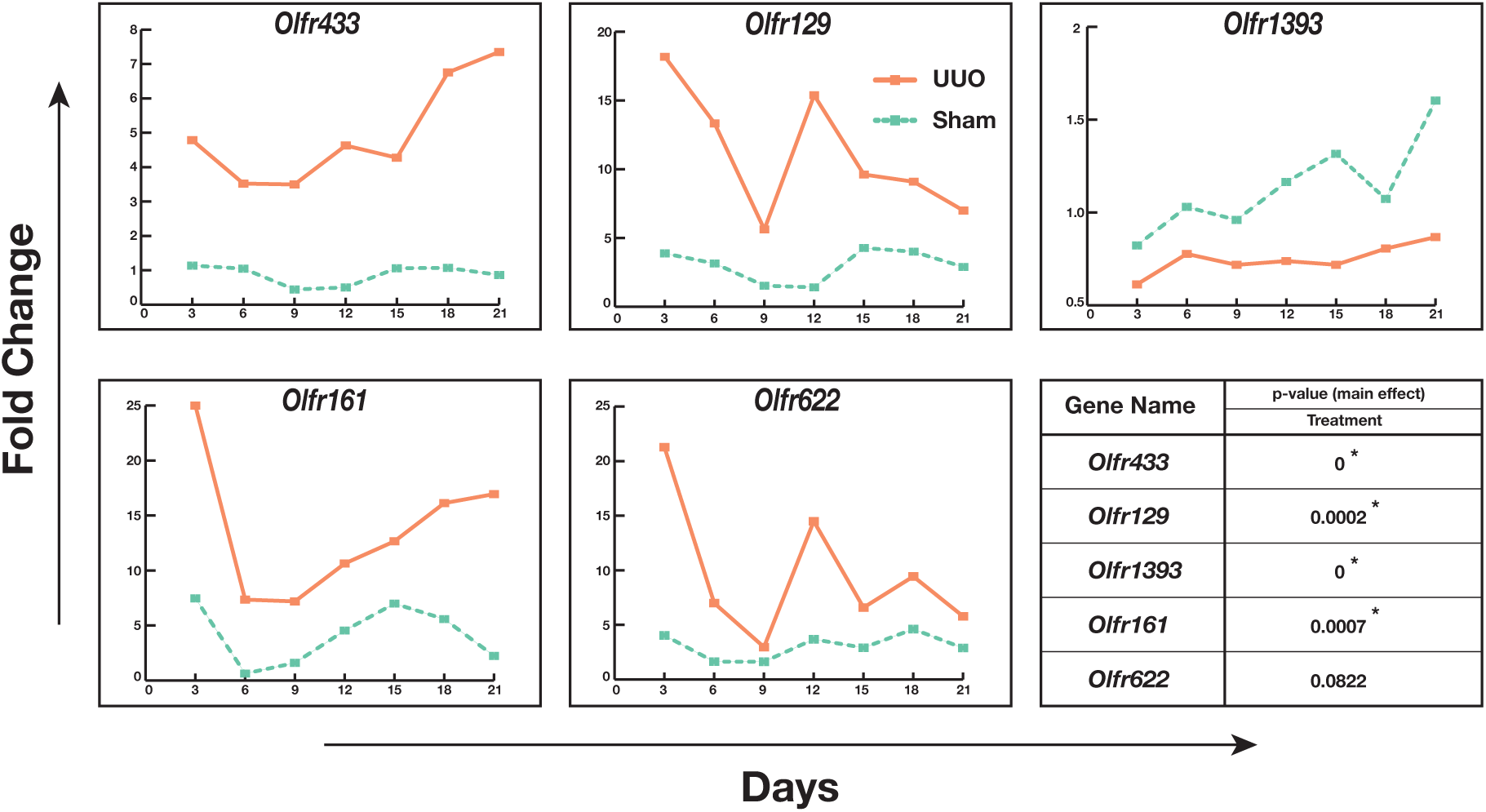
Expression level of common ORs between two datasets in mouse mode of UUO over 21 days. Statistical analysis (two-way ANOVA) determined significant changes between UUO and sham group for *Olfr433, Olfr129, Olfr1393*, and *Olfr161* (p-value < 0.05).

Considering the patterns of expression over time revealed that *Olfr433, Olfr129, Olfr161*, and *Olfr622* had a sharp downregulation from day 3 to day 6. Although *Olfr433* and *Olfr161* expression continued with a smoother augmenting response, *Olfr129* and *Olfr622* had a second sharp peak at day 12 followed by a reduction in the consecutive days. Furthermore, the expression of sham in these four genes over time revealed oscillatory patterns, which is an initial clue about the inherent rhythmic pattern of OR genes in the kidney.

## Discussion

In order to acquire a holistic view of molecular mechanisms of kidney fibrosis, two time-course microarray datasets were reanalyzed. Gene interaction map evaluation determined the modular structure of densely connected ORs in both networks. In the next step, to validate in silico results, the common ORs between two datasets were selected for further in vivo analysis in a validated model of ureteral obstruction.

Although the relationship between some of the ORs and normal physiology of the kidney has been determined in recent years (Pluznick et al., 2013; Pluznick et al., 2009; Shepard et al., 2016), few studies have focused on the role of ORs in pathological states of the kidney. The study by Shepard *et al*. has investigated the role of Olfr1393 in diabetes (Shepard et al., 2019). Additionally, our previous bioinformatics analysis determined a significant change in the expression of ORs in the rat model of Ischemia-reperfusion injury (Moradzadeh & Gheisari, 2019). To the best of our knowledge, this is the first study that has focused on the role of ORs during the progression of kidney fibrosis. Based on in silico and in vivo investigations, we identified five ORs, which according to our experimental evaluation, four of them were significantly related to the progression of kidney fibrosis in the mouse model of UUO. As these ORs have unknown ligands, deorphanizing them and functional investigations with activation-inhibition approaches would be of the matter of investigations in the future.

One of the advantages of this study is the time-course design of the models. Time-course study designs can better demonstrate the dynamism of cellular behavior and also reduces misinterpretations that may occur in single-point studies (Bar-Joseph, Gitter, & Simon, 2012; Rabieian, Moein, Khanahmad, Mortazavi, & Gheisari, 2018). Unfortunately, due to cost and difficulty, such study designs are less considered by investigators and most of the gene expression studies are performed statically (Abedi, Fatehi, Moradzadeh, & Gheisari, 2019). In the current study, both reanalyzed microarray datasets had time-series design and analyzed with a time-course analysis approach. Likewise, the in vivo model was developed in a time-course manner, both in sham and UUO groups. Following sham over time reduces the misinterpretations caused by single-time point sampling of the control group and leads to a better perspective from the dynamism of normal function.

Evaluation of *Olfr433, Olfr129, Olfr1393, Olfr161*, and *Olfr622* expression determined different patterns of gene expression over 21 days. For distinguishing real changes from noisy fluctuations, we compared the patterns of gene expression in control and treatment groups based on the parameters that we have discussed in our recent study (Rabieian et al., 2018). Accordingly, as the magnitude of difference between sham and UUO groups was significant, and the number of time points that one group was higher or lower than the other group was consistent, the expression changes seen in ORs expression were considered real and not noisy fluctuations. Furthermore, evaluations determined rhythmic patterns in ORs expression in sham groups somehow similar to the rhythmic patterns, which is not far from the oscillatory function of the kidney (Firsov & Bonny, 2018; Hara et al., 2017). Since these findings are only an initial clue about the rhythmic expression patterns of OR genes, further investigations on the relation of ORs expression patterns with kidney function would be of interest.

Taken together, our in silico and in vivo investigations validate the expression changes of novel ORs during the progression of kidney fibrosis. Our time-course evaluations determined rhythmic patterns of ORs expression and highlighted the importance of time-course study designs in deciphering dynamism of genes behavior over the course of the disease. However, future studies are required to further investigate these findings. This study is a good example of the potential capacity of systems biology unsupervised top-down strategy to unravel the neglected aspects of disease pathogenesis.

## Material and Methods

### Microarray Data analysis

The GSE36496 microarray dataset deposited by Wu *et al*. was downloaded from the gene expression omnibus (GEO) database (Barrett et al., 2013; B. Wu & Brooks, 2012). The quality of the data was assessed by Principal Component Analysis (PCA) and hierarchical clustering using ggplot2 and pheatmap package of R software, respectively (Kolde, 2019; R Core Team, 2019; Wickham, 2016). For determination of the significantly expressed genes in this time-course dataset, we applied linear models for microarray data (LIMMA), a package of R software. (Smyth, 2004). Using the multiple comparison method of LIMMA, the sham and unilateral ureteral obstruction (UUO) groups were compared with each other at different time points and differentially expressed genes (DEGs) determined by False Discovery Rate < 0.05 (Benjamini–Hochberg). The second dataset GSE96571 deposited by Wu *et al*. was also analyzed in the same way and DEGs were determined according to the previously mentioned criteria (B. Wu, Gong, Kennedy, & Brooks, 2018).

### Network Construction and topological analysis

The DEGs were used for the construction of a gene-interaction network using Cluepedia plugin (version 1.5.5) of Cytoscape software (version 3.7.2) (Bindea, Galon, & Mlecnik, 2013; Shannon et al., 2003). The interaction data for activation, binding, inhibition, and post-translational modification with a confidence rate of 0.8, was retrieved from the search tool for the retrieval of interacting genes/proteins (STRING) database (STRING-ACTION-SCORE_v10.0_10090_09.06.2015) (Szklarczyk et al., 2017). Based on the clustering coefficient parameter, the densely connected sites of the network with a cutoff point of 4 were determined by molecular complex detection (MCODE) plugin (version 1.5.1) as structural modules (Bader & Hogue, 2003). The gene-interaction network for GSE96571 microarray dataset was also constructed, and the structural modules were investigated using the same method.

### Animal model of Unilateral Ureteral Obstruction

Male C57BL/6 mice aged 6-8 weeks were obtained from Pasteur Institute (Tehran, Iran). Animal care and experiments were according to the National Institutes of Health’s Guide for the Care and Use of Laboratory Animals. Ketamine and Xylazine (Alfasan, Woerden, Netherland) were injected intraperitoneally for anesthesia at the dose of 115 and 11.5 mg/kg, respectively. During anesthesia and operation, mice were kept on a 37.5 °C plate. After a mid-abdominal incision, the left ureter was isolated from surrounding tissues, double ligated, and the incision was sutured. Sham operation was performed with the same procedure, except for the ligation of the ureter. The mice were harvested 1, 3, 6, 9, 12, 15, 18, and 21 days after surgery. For each time point, two UUO and two sham-operated mice were allocated. Besides, four untreated mice were used as normal controls. After sacrificing with cervical dislocation, the left kidney was harvested and coronal sections were prepared. The anterior part was kept in 3.7% formaldehyde in PBS for histopathological study and the posterior part was sustained in liquid nitrogen for RNA extraction.

### Histopathological Evaluation

The formalin-fixed kidney tissues were paraffin-embedded and 5µm sections were prepared. Hematoxylin and eosin (H&E) as well as Masson trichrome staining were carried out and histopathological evaluations were performed in a blinded manner. In random cortical fields, the existence of increased mesangial matrix, diffused glomerulosclerosis, and mesangial cell proliferation was assessed between two experimental groups. Moreover, for each section, the percentage of cortex area affected by fibrosis that included interstitial cortical fibrosis, tubular loss with minimal fibrosis, and periglomerular fibrosis was determined as described previously (Moghadasali et al., 2015).

### Statistical Analysis

To compare the significance of histopathological differences between sham and UUO groups, the two-way ANOVA test was applied using the ‘anova2’ function of MATLAB software (MathWorks, 2019a).

### Gene Expression Assessment

Total RNA of the lower part of the left kidney was extracted using RNX-plus (CinnaGen, Tehran, Iran) according to the manufacturer’s instruction. Afterward, the concentration of the extracted RNA was measured by Epoch microplate spectrophotometer (BioTek, Winooski, Vermont). Since ORs sequence contains only the exon coding region, to eliminate DNA contamination, the samples were treated with DNase I (Thermo Fisher, Waltham, Massachusetts). To validate this procedure, mock controls were also employed. Subsequently, random hexamer primered cDNA synthesis was done using the first-strand cDNA synthesis kit (YektaTajhiz, Tehran, Iran). Specific primers for the selected ORs, *Hprt* and *Tfrc* were designed using AlleleID software version 6.2 (Supplementary Table 1) (Apte & Singh, 2007). Real-time quantitative polymerase chain reaction (RT-qPCR) was performed using RealQ Plus 2x Master Mix Green with high ROX™ (Ampliqon, Odense, Denmark) by Rotor-gene 6000 cycler (Qiagen, Hilden, Germany). The expression of genes was normalized by considering *Hprt* and *Tfrc* as internal control genes. The results of RT-qPCR were analyzed using the relative expression software tool (REST) version 1 (Pfaffl, Horgan, & Dempfle, 2002).

### Statistical Analysis

To compare gene expression patterns in UUO and sham groups, the two-way ANOVA test was applied using the ‘anova2’ function of MATLAB software (MathWorks, 2019a). The fold changes of the UUO group were considered as response values.

### PCR products sequencing

PCR was performed by T100™ Thermal Cycler (Bio-Rad, Hercules, California) using Taq DNA Polymerase 2x Master Mix RED (Ampliqon, Odense, Denmark), then the PCR products were cloned into the PTZ57R vector using TA Cloning™ kit (ThermoFisher, Waltham, Massachusetts). Cloned products were transformed into competent TOP10 *E. coli* by incubating on ice with a subsequent heat-shock at 37°C. Transformed colonies were cultured on LB-Agar (Sigma-Aldrich, St. Louis, Missouri) plate treated with ampicillin followed by overnight incubation at 37°C. Afterward, plasmid extraction from bacteria was performed using Solg™ Plasmid Mini-prep Kit (SolGent, Daejeon, South Korea). Sanger sequencing of samples was done using both forward and reverse universal M13 (−40) primers by Bioneer biotech company (Daejeon, South Korea).

## Acknowledgments

This study was supported by Isfahan University of Medical Sciences (grant number: 398439).

---

**Supplementary Table 1:**
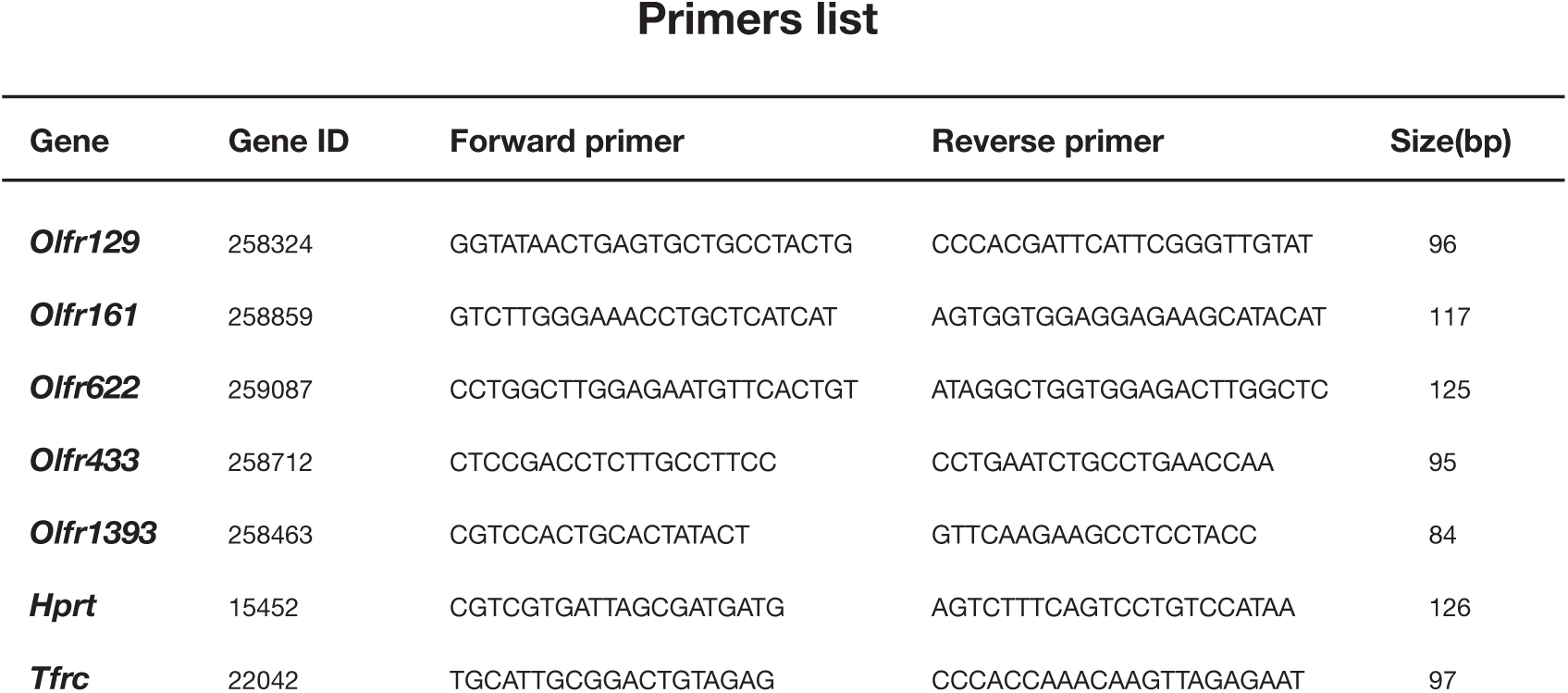
Supplementary Data 1: Sanger sequencing results

